## Supplemental Material for "A marine cryptochrome with an inverse photo-oligomerization mechanism"

### Contents

Supplementary Figure 1

Supplementary Figure 2

Supplementary Figure 3

Supplementary Figure 4

Supplementary Figure 5

Supplementary Figure 6

Supplementary Figure 7

Supplementary Table 1

Supplementary Table 2

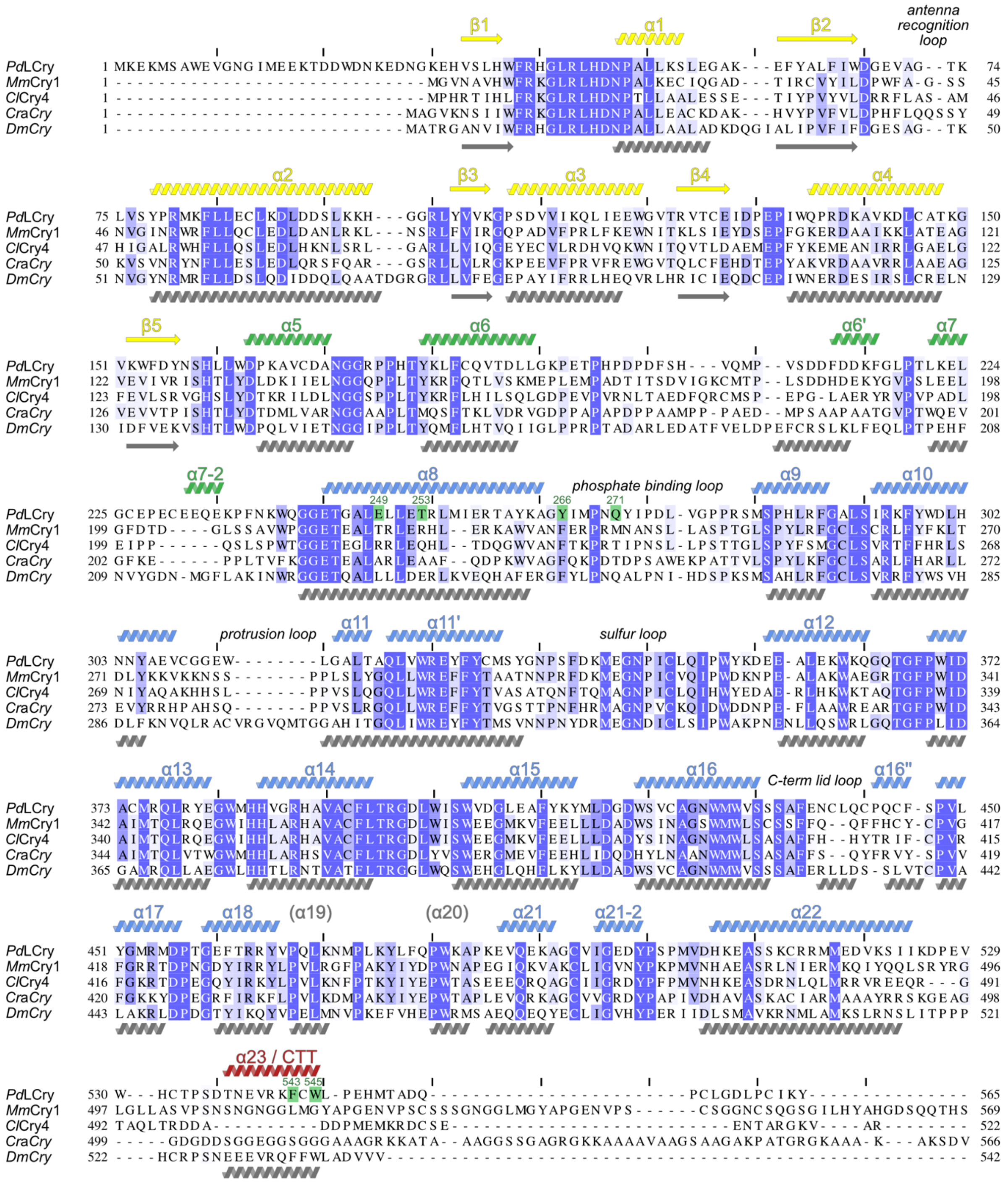


**Supplementary Figure 1: Sequence alignment of animal- and animal-like CRYs with known structure.**

Detailed sequence alignment of *Pd*LCry (NCBI AccNo. AGX93007), mouse cryptochrome 1 (*Mm*CRY1: P97784), pigeon cryptochrome 4 (*Cl*Cry4: ANT47196), green algae animal-like cryptochrome (*Cra*Cry: A8J8W0) and drosophila cryptochrome (*Dm*Cry: NP_732407). Secondary structure elements of *Pd*LCry (coloured according to the domain organisation of Fig. 1, based on our dark state structure) and *Dm*Cry (grey) are shown. For *Dm*Cry, these elements and their numbering are based on Czarna *et al.* (*Czarna, A. et al. Structures of Drosophila Cryptochrome and Mouse Cryptochrome1 Provide Insight into Circadian Function. Cell 153, 1394–1405 (2013)*). Amino acids mutated in this study are highlighted in green.

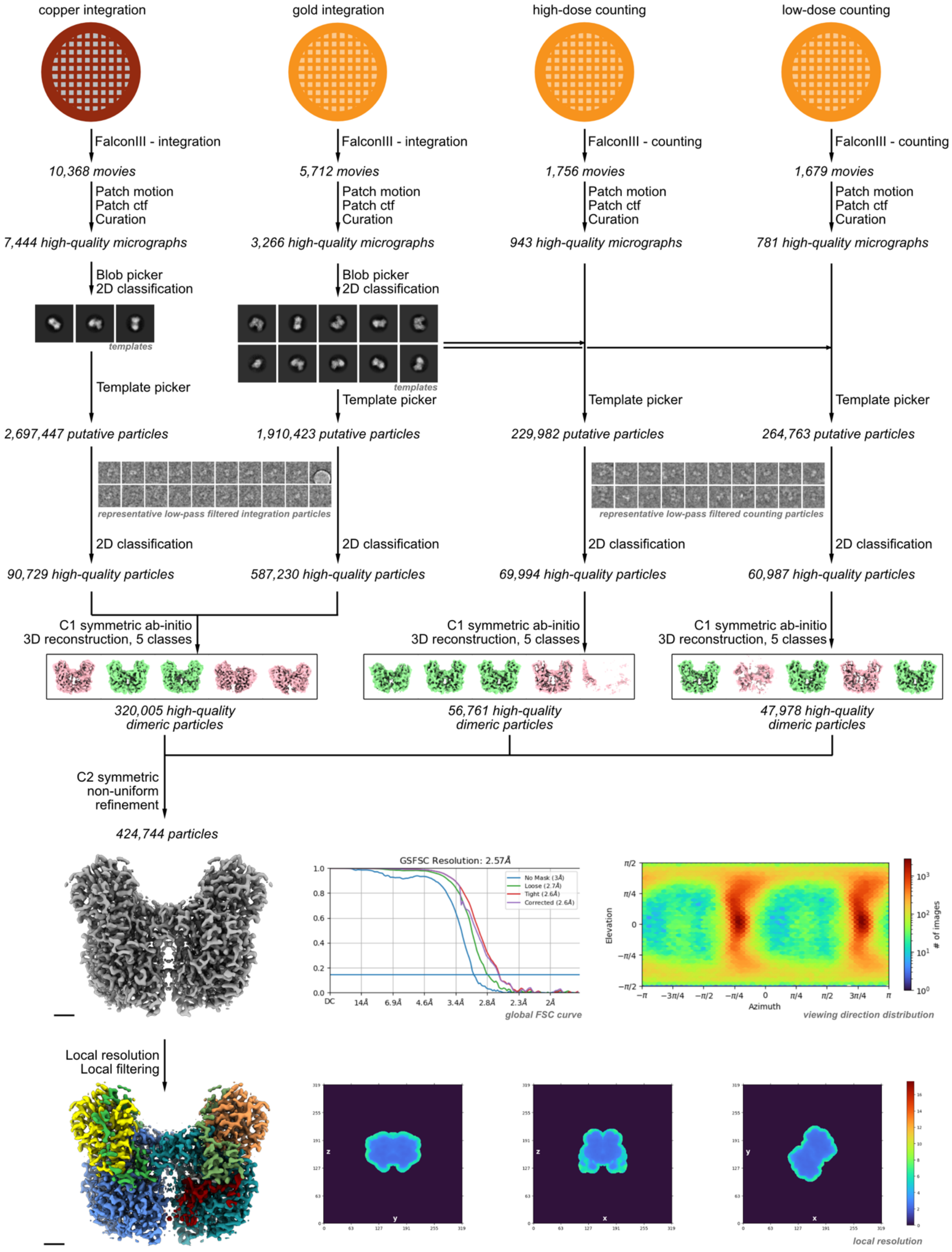


**Supplementary Figure 2. Overview of the single particle cryo-EM data processing workflow for the dark state.**

A total of 4 datasets were collected on a 300 kV cryo-electron microscope. Two of these datasets were acquired in the integration mode of the Falcon III detector, and the other two were acquired in the counting mode. Movies were selected for low per-frame drift rates, good CTF scores, and low astigmatism. Particles were picked using a combination of blob and template picker, and then curated using unsupervised 2D classification, selecting for particles with protein-like density and resolutions better than 7 Å. This resulted in a set of both monomeric (about 15%) and dimeric (about 80%) particles. The dimeric particles were further curated using ab-initio reconstruction, sorting them into 5 distinct populations. From these all particles contributing to C2-symmetric dimeric structures (shown in green) were combined and refined in 3D using a non-uniform refinement algorithm, resulting in a map with a uniform resolution of 2.6Å. For further details, see the Materials and Methods, and Supplementary Table 2.

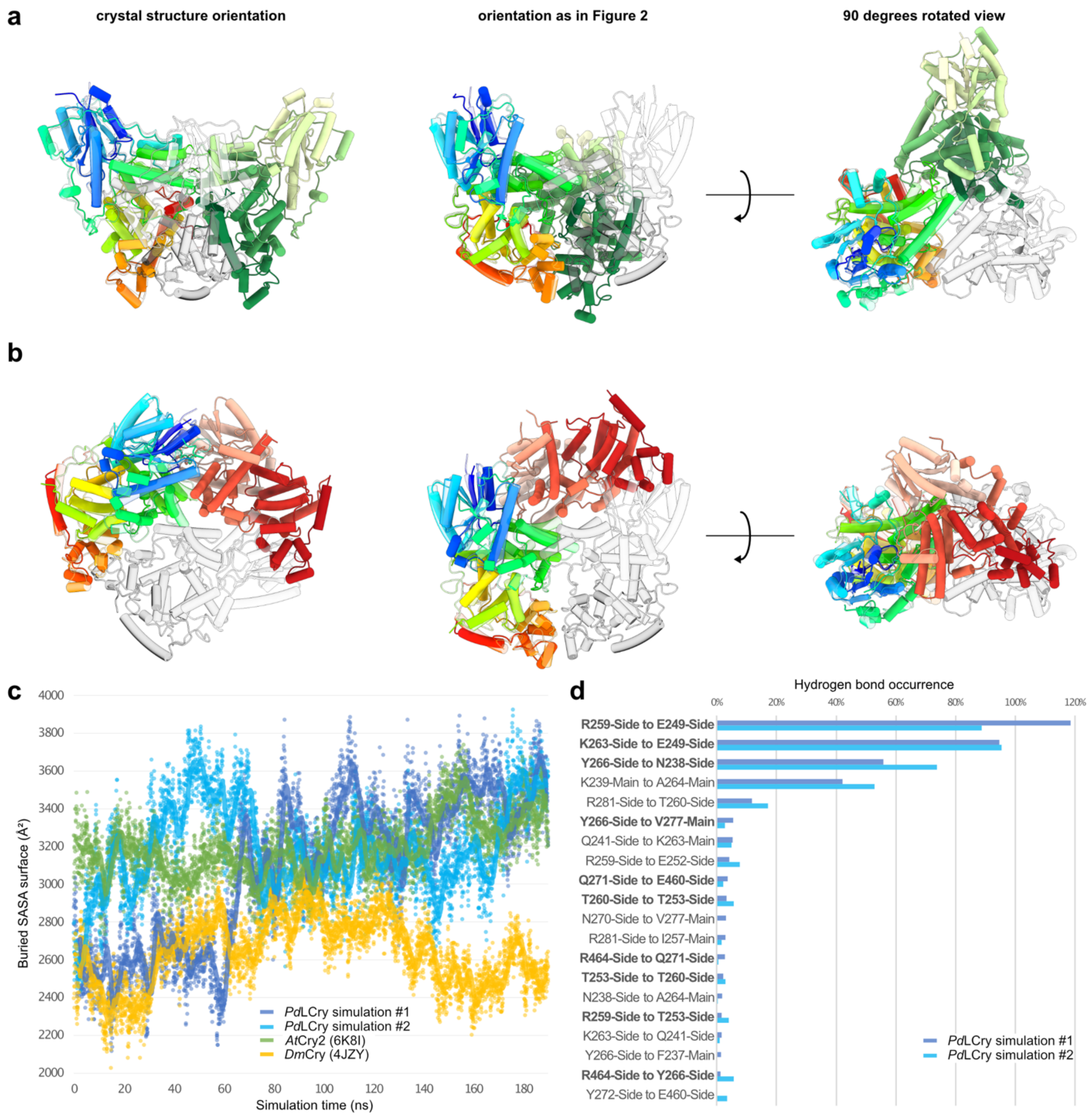


**Supplementary Figure 3: Comparison of the *Pd*LCry dark state dimer with other CRY dimer arrangements.**

**a** Comparison of the subunit arrangement in the *Dm*Cry crystal dimer and **b** in the dark state *At*Cry2 dimer with the arrangement in the dark state *Pd*LCry dimer (shown as transparent). For each, subunit 1 has been aligned to subunit 1 of the *Pd*LCry dimer. These subunits are depicted using a rainbow color gradient from blue (N-terminus) to red (C-terminus). Subunit 2 is depicted in a gradient from light green (N-terminus) to dark green (C-terminus) for *Dm*Cry (4JZY), from light red (N-terminus) to dark red (C-terminus) for *At*Cry2 (6K8I), and from transparent light gray (N-terminus) to transparent dark gray (C-terminus) for *Pd*LCry. **c** Evolution of the solvent accessible surface area (SASA) buried in the interaction interface during MD simulations of the *Pd*LCry dimer (two independent simulations, light and dark blue traces), the physiologically relevant *At*Cry2 dimer (green trace), and the crystal *Dm*Cry dimer (orange trace). Considering the first 190 ns of the simulation, *At*Cry2 dimers (SASA average 3194 ± 164 Å$^2$) and *Pd*LCry dimers (SASA average 3138 ± 343 Å$^2$) had on average larger buried interfaces compared to the unphysiological *Dm*Cry dimer (SASA average 2640 ± 189 Å$^2$). Individual values at each time point of the simulation are shown as transparent dots, the lines show the running average over 0.25 ns simulation time. **d** Number of dimer-interface hydrogen bonds observed during the two independent *Pd*LCry MD simulations, with interactions involving residues that were studied in follow-up mutagenesis experiments (see Fig 6) shown in bold. The list is limited to hydrogen bonds involving residues of helix α8 and its flanking region (225 – 302), and of helix α18 (459 – 465), see Fig. 3. Since several sidechains can form more than one hydrogen bond simultaneously, occurrences above 100% are possible (e.g. Arg259 to Glu249 interaction).

**a**

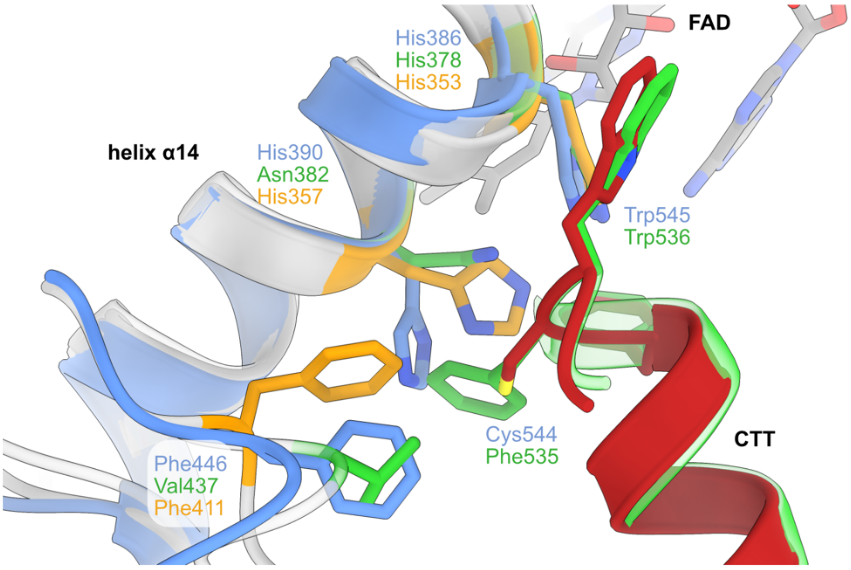


**b**

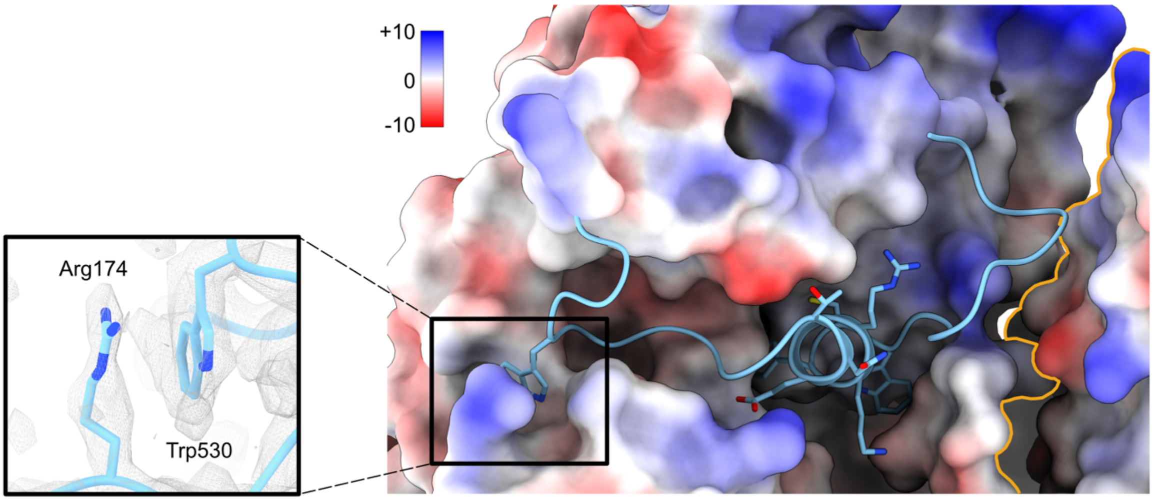


**Supplementary Figure 4: Details of the CTE binding surface.**

**a** Comparison of *Pd*LCry (blue, CTT red), *Dm*Cry (green, PDB 6WTB), and *Cl*Cry4 (orange, PDB 6PU0) interactions with the CTT. Note that the crystallization construct used for solving the structure of *Cl*Cry4 does not contain a CTT, and that our sequence alignment does not show an FFW/FCW-related motif in *Cl*Cry4 (Supplementary Fig. 1). While His386 of *Pd*LCry, which bridges between the CTT and the FAD chromophore, is conserved in all three CRYs, His390 and Phe446 are present only in *Pd*LCry and the type IV CRY *Cl*Cry4. However, *Cl*Cry4 His357 is in a rotamer position reminiscent of *Dm*Cry Asn382, possibly due to the truncation of the CTT in the crystallization construct. *Dm*Cry Phe535 would clash with His390 of *Pd*LCry. The different position of *Cl*Cry4 Phe411 compared to *Pd*LCry Phe446 could also be due to the absence of a CTT in the truncated crystallization construct. **b** Surface representation of the *Pd*LCry atomic model colored by its electrostatic potential ranging from +10 to -10 kcal/(mol·*e*). The *Pd*LCry CTE (blue ribbon) has an additional anchor N-terminal to the CTT, namely Trp530, which interacts with Arg174 (see insert). The orange line delineates subunit 1 from subunit 2.

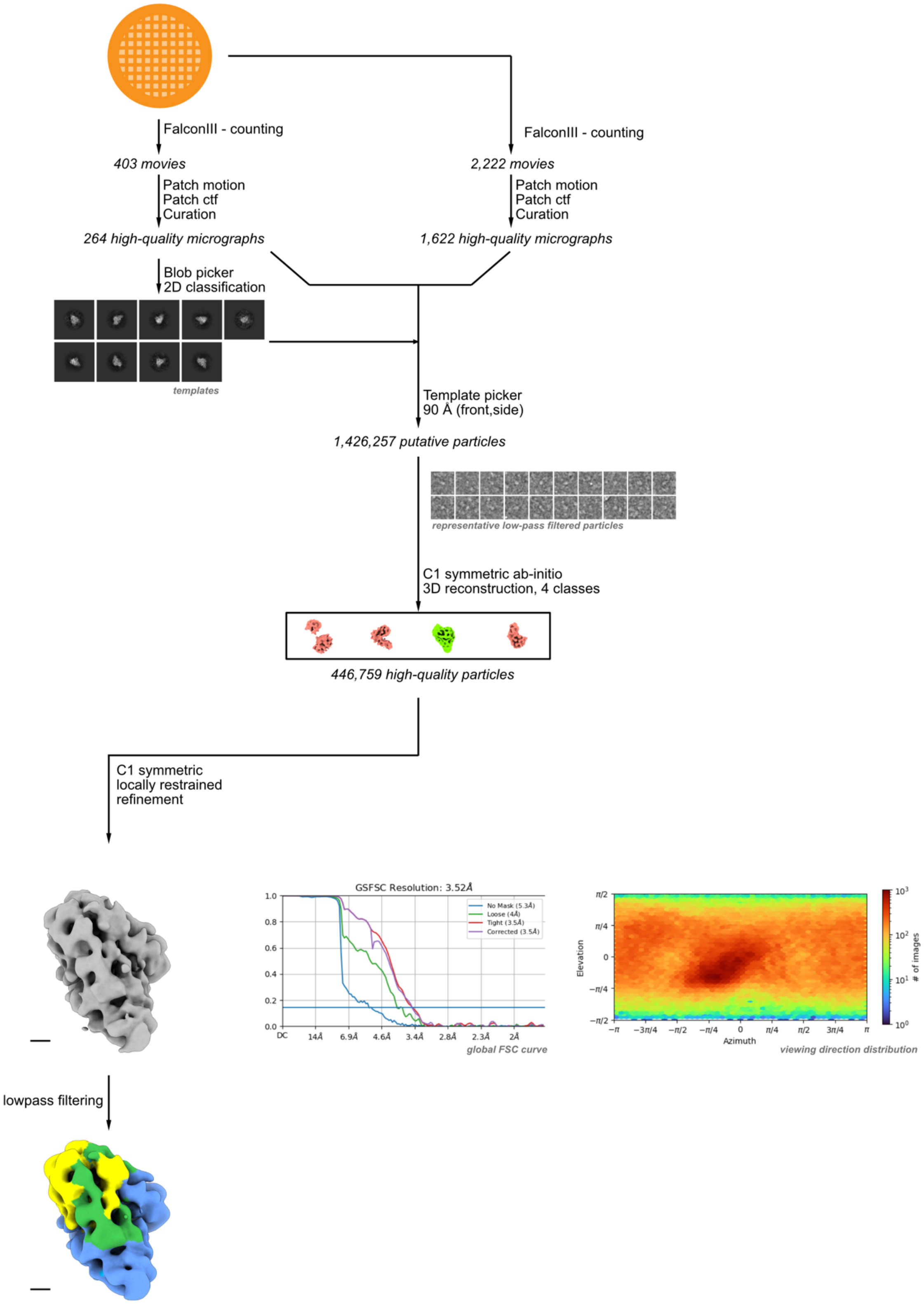


**Supplementary Figure 5. Overview of the single particle cryo-EM data processing workflow for the blue light state.**

A total of 2 data sets were collected on a 300 kV cryo-electron microscope in the counting mode of the Falcon III detector. Movies were selected for low per-frame drift rates, good CTF scores and low astigmatism. Particles were picked using a combination of blob and template picker. Unsupervised 2D classification revealed that most of them (91%) were clearly monomeric. For further refinement, all particles were classified in 3D using ab-initio reconstruction, sorting into 4 distinct populations. Of these, only one showed a continuous, protein-like density and particles contributing to this subset were refined in 3D using a non-uniform refinement algorithm with local priors to stabilize the reconstruction, yielding a map with a nominal resolution of 3.5Å, but which was anisotropic. For interpretation we low-pass filtered this map to 8 Å. For further details, see the Materials and Methods section and Supplementary Table 2.

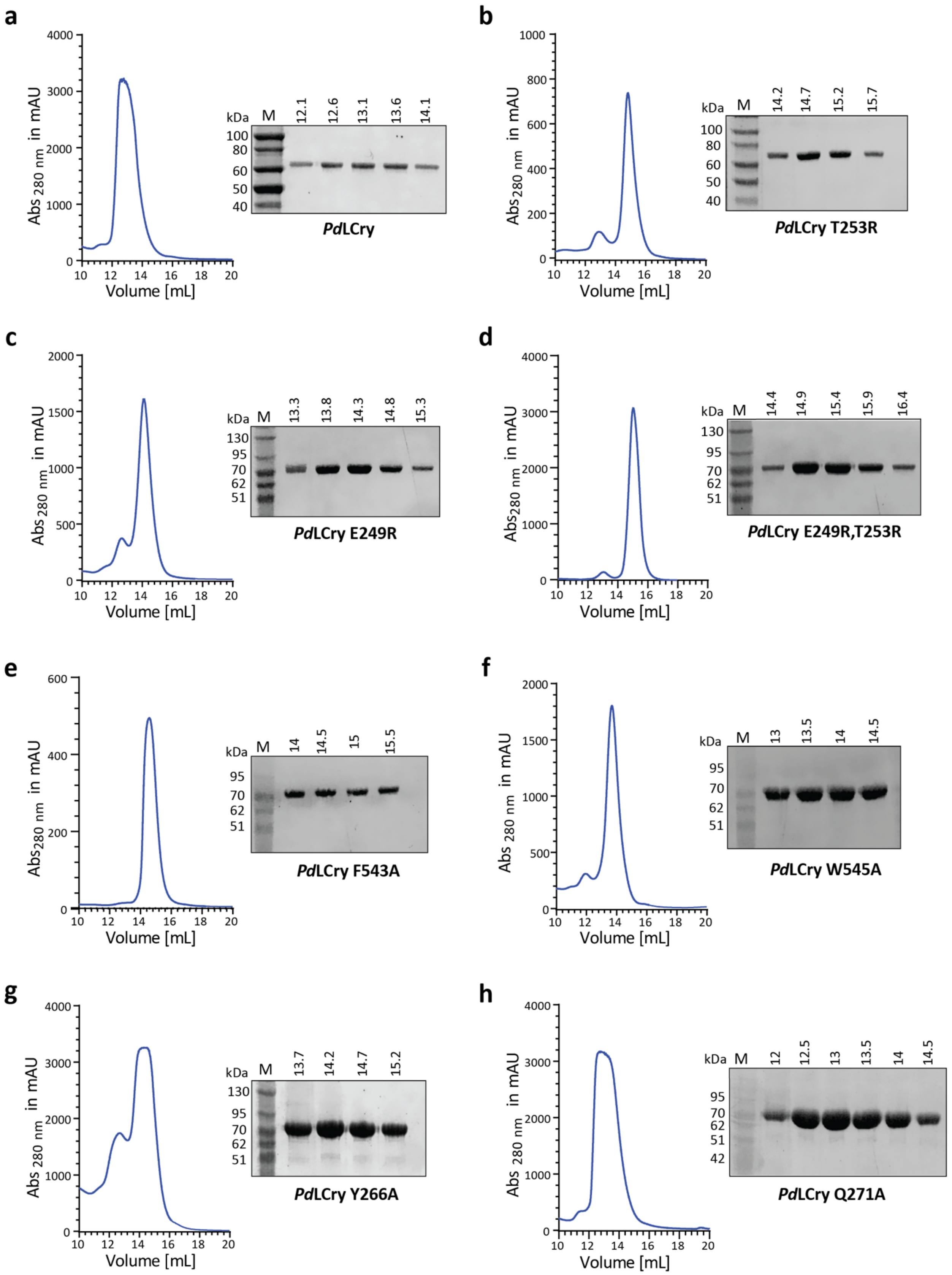


**Supplementary Figure 6: Final SEC purification step of wildtype and mutant *Pd*LCry proteins.**

Size exclusion chromatography (SEC) chromatograms using an S200 10/300 column (left) and subsequent SDS-PAGE analyses of the SEC peak fractions (right) of **a** wildtype *Pd*LCry, **b** *Pd*LCry$^{T253R}$, **c** *Pd*LCry$^{E249R}$, **d** *Pd*LCry$^{E249R,T253R}$, **e** *Pd*LCry$^{F543A}$, **f** *Pd*LCry$^{W545A}$, **g** *Pd*LCry$^{Y266A}$ and **h** *Pd*LCry$^{Q271A}$. Elution volumes (in mL) of each SEC fraction are indicated on top of the SDS-PAGE gels. M: MW marker. Total SEC peaks (wt *Pd*LCry and *Pd*LCry$^{Q271A}$) or monomer peaks (all mutants except *Pd*LCry$^{Q271A}$) were pooled and concentrated for the further analyses.

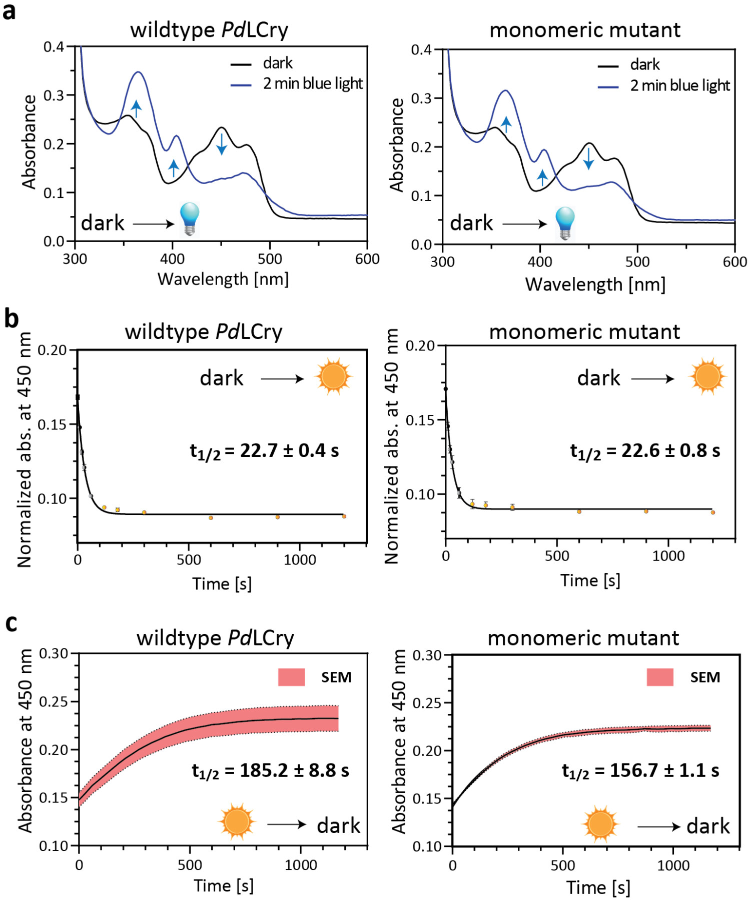


**d**

| | wildtype $t_{1/2}$ [s] | monomer $t_{1/2}$ [s] |
|---|---|---|
| Sunlight photoreduction (4°C) | 22.1 | 21.6 |
| | 22.7 | 24.3 |
| | 23.4 | 22.1 |
| Sunlight dark recovery (18°C) | 137.1 | 116.4 |
| | 151 | 122.1 |
| | 201 | 169.6 |

**Supplementary Figure 7: Blue light and sunlight photoreduction and dark recovery kinetics of wildtype *Pd*LCry and the monomeric *Pd*LCry E249R T253R mutant.**

**a** Absorption spectra of wildtype *(left)* and mutant *(right)* *Pd*LCry in darkness (black) and after 2 min blue light exposure (blue). Arrows indicate the change in absorbance at 370 nm, 404 nm and 450 nm upon photoreduction of $FAD^{OX}$ (black) to the anionic $FAD^{ASQ}$ radical (blue). **b** Sunlight photoreduction kinetics of wildtype *(left)* and mutant *(right)* *Pd*LCry following absorbance changes at 450 nm (on ice, ≡ 4°C). Normalized average data points obtained from three independent measurements are plotted. Error bars represent SEM of the three independent replicates. The $t_{1/2}$ values are means of three replicates +/- SEM. **c** Dark recovery kinetics of wildtype *(left)* and mutant *(right)* *Pd*LCry after sunlight, following absorbance changes at 450 nm at 18°C. Average data points obtained from three independent measurements are plotted. The red shaded region represents the standard error of the mean (SEM) of three independent replicates (at each data point). The $t_{1/2}$ values are means of three replicates +/- SEM. **d** Individual half-live ($t_{1/2}$) values of three independent measurements of sunlight photoreduction and dark recovery kinetics for wildtype *Pd*LCry and the monomeric E249R T253R mutant.

**Supplementary Table 1: Sequence and species information**

| name in figure 1 | protein name | species | NCBI/Uniprot ID | comment |
|---|---|---|---|---|
| AgCry1 | cryptochrome 1 | *Anopheles gambiae* | ABB29886.1 | |
| AgCry2 | cryptochrome 2 | *Anopheles gambiae* | ABB29887.1 | |
| AmCry2 | cryptochrome 2 | *Apis mellifera* | NP_001077099.1 | |
| AquCry1 | cryptochrome-2-like | *Amphimedon queenslandica* | XP_003386582.1 | |
| AquCry2 | cryptochrome-2-like | *Amphimedon queenslandica* | XP_003386569.1 | |
| AtCry1 | cryptochrome 1 | *Arabidopsis thaliana* | NP_567341.1 | |
| AtCry2 | cryptochrome 2 | *Arabidopsis thaliana* | NP_849588.1 | |
| AtCry3 | cryptochrome 3 | *Arabidopsis thaliana* | NP_568461.3 | |
| ClCry1 | cryptochrome 1 | *Columba livia* | NP_001302444.1 | |
| ClCry4 | cryptochrome 4 | *Columba livia* | ANT47196.1 | |
| CmeCry | cryptochrome | *Crateromorpha meyeri* | CAZ66368.1 | |
| CraCry | Uncharacterized protein | *Chlamydomonas reinhardtii* | A8J8W0 | from PDB 5ZMO |
| DmCry | cryptochrome | *Drosophila melanogaster* | NP_732407.1 | |
| DpCry1 | cryptochrome 1 | *Danaus plexippus* | XP_032522602.1 | |
| DpCry2 | cryptochrome 2 | *Danaus plexippus* | ABA62409.1 | |
| DrCry3 | cryptochrome 3 | *Danio rerio* | BAA96850.1 | |
| DrCry4 | cryptochrome circadian regulator 4 | *Danio rerio* | NP_571862.1 | |
| DrCryDASH | cryptochrome DASH | *Danio rerio* | NP_991249.1 | |
| ErCry1 | cryptochrome 1 | *Erithacus rubecula* | Q5IZC5.2 | |
| ErCry4 | cryptochrome 4 | *Erithacus rubecula* | ATE87950.1 | |
| GgCry1 | cryptochrome 1 | *Gallus gallus* | NP_989576.1 | |
| GgCry2 | cryptochrome 2 | *Gallus gallus* | NP_989575.1 | |
| GgCry4 | cryptochrome 4 | *Gallus gallus* | NP_001034685.1 | |
| GviCryDASH | DASH family cryptochrome | *Gloeobacter violaceus* | WP_011140837.1 | |
| HsCry1 | cryptochrome 1 | *Homo sapiens* | NP_004066.1 | |
| HsCry2 | cryptochrome 2 | *Homo sapiens* | NP_066940.3 | |
| MmCry1 | cryptochrome 1 | *Mus musculus* | P97784.1 | |
| MmCry2 | cryptochrome 2 | *Mus musculus* | Q9R194.1 | |
| PdLCry | light receptive cryptochrome | *Platynereis dumerilii* | UUF95169 | |
| SdoCry | cryptochrome | *Suberites domuncula* | CAZ66367.1 | |
| SeCry | cryptochrome | *Spodoptera exigua* | ADY17887.1 | |
| SlCry1 | cryptochrome 1 | *Solanum lycopersicum* | NP_001234667.1 | |
| SlCry2 | cryptochrome 2 | *Solanum lycopersicum* | NP_001234245.1 | |
| SlCryDASH | cryptochrome DASH | *Solanum lycopersicum* | NP_001296304.1 | |
| VcCryDASH | DASH family cryptochrome | *Vibrio cholerae* | WP_000037642.1 | |
| XlCry2 | cryptochrome circadian regulator 2 S homeolog | *Xenopus laevis* | NP_001083936.1 | |
| XlCryDASH | cryptochrome DASH | *Xenopus laevis* | NP_001084438.1 | |
| XtCry4 | cryptochrome 4 isoform X1 | *Xenopus tropicalis* | XP_012812443.1 | |

**Supplementary Table 2: Statistics and additional information of the refinement and model building process**

| **Dataset dark state** | Cu integration | Au integration | high-dose counting | low-dose counting |
|---|---|---|---|---|
| **Sample conditions** | | | | |
| Sample concentration | 0.7 mg/ml | | | |
| Additive | 0.01% fOM | | | |
| Grid type | Quantifoil Cu R1.2/1.3. (300 mesh) | UltrAUfoil R1.2/1.3 (300 mesh) | | |
| Light condition | Far-red (>680 nm) | | | |
| **Cryo-EM data collection** | | | | |
| Microscope | Titan Krios G3i | | | |
| Voltage (kV) | 300 | | | |
| Spherical aberration Cs (mm) | 2.7 | | | |
| Condenser C2 aperture size (µm) | 100 | 100 | 50 | 50 |
| Objective aperture size (µm) | No aperture | No aperture | 100 | 100 |
| Camera | Falcon III | | | |
| Pixel size (Å) | 0.862 | | | |
| Total dose (electron*Å$^{-2}$) | 66 | | 27 | |
| Number of frames | 19 | | 24 | 48 |
| Exposure time (sec) | 0.5 | | 39.5 | 39.5 |
| Images per hole | 2 | 1 | 2 | 2 |
| Energy filter | None | | | |
| Defocus range (µm) | -2.0 to -1.0 | -1.9 to -1.0 | -2.0 to -0.3 | -2.0 to -0.3 |
| # micrographs collected | 10,368 | 5,712 | 1,756 | 1,679 |
| % micrographs used | 72 | 57 | 54 | 47 |
| **Cryo-EM data processing** | | | | |
| Software | cryoSPARC v3.3.2+220518 patch | | | |
| Picked particles | 2,697,447 | 1,910,423 | 229,982 | 264,763 |
| Particles after 2D classification | 90,729 | 587,230 | 69,994 | 60,987 |
| Particles after 3D sorting | 320,005 | | 56,761 | 47,978 |
| Resolution (FSC 0.143, Å) | 2.6 | | | |
| **Model building and refinement** | | | | |
| Software for building | Coot 0.9.4.7 EL | | | |
| Residues build | 31-556 | | | |
| Software for refinement | PHENIX 1.20.1 - 4487 | | | |
| Composition (#)<br>Chains<br>Atoms<br>Residues<br>Water<br>Ligands | <br>2<br>8756 (Hydrogens: 0)<br>Protein: 1052<br>84<br>Mg: 2, FAD: 2 | | | |
| Bonds (RMSD)<br>Length (Å) (# > 4$\sigma$)<br>Angles (°) (# > 4$\sigma$) | <br>0.003<br>0.511 | | | |
| Ramachandran plot (%)<br>Outliers<br>Allowed<br>Favored | <br>0<br>3.63<br>96.37 | | | |
| Rotamer Outliers (%) | 0.0 | | | |
| MolProbity score | 1.15 | | | |

| **Dataset blue light state** | counting |
|---|---|
| **Sample conditions** | |
| Sample concentration | 0.7 mg/ml |
| Additive | none |
| Grid type | UltrAUfoil<br>R1.2/1.3<br>(300 mesh) |
| Light condition | background illumination: Far-red (>680 nm); sample: 30 sec 455 nm blue LED |
| **Cryo-EM data collection** | |
| Microscope | Titan Krios G3i |
| Voltage (kV) | 300 |
| Spherical aberration Cs (mm) | 2.7 |
| Condenser C2 aperture size (µm) | 70 |
| Objective aperture size (µm) | 100 |
| Camera | Falcon III |
| Pixel size (Å) | 0.862 |
| Total dose (electron*$Å^{-2}$) | 30 |
| Number of frames | 36 |
| Exposure time (sec) | 54 |
| Images per hole | 1 |
| Energy filter | None |
| Defocus range (µm) | -2.0 to -0.3 |
| # micrographs collected | 2,625 |
| % micrographs used | 72 |
| **Cryo-EM data processing** | |
| Software | cryoSPARC v3.3.2+220518 patch |
| Picked particles | 1,426,257 |
| Particles after 3D sorting | 446,759 |
| Resolution (FSC 0.143, Å) | 3.5 (filtered to 8 Å) |
